# Choice of 16S ribosomal RNA primers impacts urinary microbiota profiling

**DOI:** 10.1101/2022.01.24.477608

**Authors:** Vitor Heidrich, Lilian T. Inoue, Paula F. Asprino, Fabiana Bettoni, Antonio C.H. Mariotti, Diogo A. Bastos, Denis L.F. Jardim, Marco A. Arap, Anamaria A. Camargo

## Abstract

Accessibility to next-generation sequencing (NGS) technologies has enabled the profiling of microbial communities living in distinct habitats. 16S ribosomal RNA (rRNA) gene sequencing is widely used for microbiota profiling with NGS technologies. Since most used NGS platforms generate short reads, sequencing the full-length 16S rRNA gene is impractical. Therefore, choosing which 16S rRNA hypervariable region to sequence is critical in microbiota profiling studies. All nine 16S rRNA hypervariable regions are taxonomically informative, but due to variability in profiling performance for specific clades, choosing the ideal 16S rRNA hypervariable region will depend on the bacterial composition of the habitat under study. Recently, NGS allowed the identification of microbes in the urinary tract, and urinary microbiota has become an active research area. However, there is no current study evaluating the performance of different 16S rRNA hypervariable regions for urinary microbiota profiling. We collected urine samples from male volunteers and profiled their urinary microbiota by sequencing a panel of six amplicons encompassing all nine 16S rRNA hypervariable regions. After systematically comparing their performance, we show that V1V2 hypervariable regions better assess the taxa commonly present in urine samples and V1V2 amplicon sequencing is more suitable for urinary microbiota profiling. We believe our results will be helpful to guide this crucial methodological choice in future urinary microbiota studies.

## Introduction

Urine is not sterile (1). Modified culture protocols and modern sequencing techniques have now enabled the detection of microbes washed out from the whole urogenital tract (2). However, only a small fraction of these microbes is culturable (3), rendering culture-independent sequencing-based methods as the main tool to identify microbes inhabiting the urogenital tract.

Microbial communities colonizing the urinary tract (collectively referred to as the urobiome) are influenced by sex, age, environmental factors and even host genetics (2,4). Most importantly, recent studies have shown that urobiome dysbiosis is linked to several urological conditions (2), ranging from urinary incontinence (5) to bladder cancer (6). Therefore, a comprehensive and systematic characterization of the urobiome in health and disease is fundamental, and may lead to new prevention, diagnosis and treatment strategies for urological pathologies.

Bacteria are the dominant component of the urobiome and a major technical challenge in DNA-based microbiota studies is the low bacterial biomass of urine samples. The urinary tract contains <10^5^ colony forming units per milliliter, a number at least a million times lower than that found in feces per gram (7). As a consequence, while gut microbiota DNA-based studies are shifting from 16S ribosomal RNA (rRNA) amplicon sequencing towards shotgun metagenomic sequencing - which is problematic with low amounts of input bacterial DNA (8) -, urinary microbiota profiling still relies on 16S rRNA amplicon sequencing (9).

A critical step in 16S rRNA amplicon sequencing studies is the selection of which 16S rRNA hypervariable regions to sequence. 16S rRNA contains nine hypervariable regions (V1-V9) used to determine taxonomic identity and estimate evolutionary relationships between bacteria. Although all nine hypervariable regions are taxonomically informative, the amount and quality of information retrieved varies per region according to the studied environment. For instance, Fadeev et al. (2021) showed that V4V5 is superior to V3V4 for microbiota profiling of environmental arctic samples (10), and Kameoka et al. (2021) found that V1V2 is more precise than V3V4 for gut microbiota profiling of Japanese individuals (11).

Despite evidence showing that the choice of 16S hypervariable regions in microbiota profiling studies is critical (10–13), no study has systematically compared the performance of different 16S rRNA hypervariable regions for microbial characterization of urine samples. In this work, we compared the performance of different sets of 16S rRNA primers for urinary microbiota profiling. We collected urine samples from male volunteers by transurethral catheterization and used a 16S rRNA sequencing panel encompassing all nine hypervariable regions. We also combined pairs of non-overlapping 16S rRNA amplicons using bioinformatics reconstruction to evaluate their performance. To identify which primer sets and combinations are best suited for urinary microbiota profiling, we evaluated the effect of using different primer sets and combinations on metrics such as taxonomic resolution, taxonomic richness and ambiguity. We show that V1V2 amplicon sequencing is more suitable for urinary microbiota studies. We also observed marginal gains in taxonomic richness when using pairs of amplicons, which may not compensate for the higher costs of sequencing multi-amplicon libraries.

## Materials and Methods

### Sample collection

Twenty-two urine samples were collected from 14 male volunteers between March 2019 and November 2020. Samples were collected by a trained nurse in sterile urine containers during catheterization for BCG instillation in volunteers with non-muscle invasive bladder cancer or for transurethral resection in volunteers with benign prostatic hyperplasia (Table S1). Urine samples were stored at −80 °C until DNA extraction.

### DNA extraction

Urine samples were thawed at room temperature, and up to 40 ml of urine was used for DNA extraction. Urine samples were centrifuged twice for 15 min at 10 °C and 3000 g, and the supernatant was discarded sparing 10 ml of urine (containing a pellet). This content was transferred to 15 ml tubes, and centrifugation was repeated (15 min; 10 °C; 3000 g). Approximately 1 ml of urine (containing the pellet) was resuspended in 3 ml phosphate-buffered saline (PBS) and centrifugation was repeated (15 min; 10 °C; 3000 g). The supernatant was discarded leaving 1 ml of sample in the tube. Samples and the DNA extraction negative control (1 ml PBS) were processed for DNA extraction using the QIAamp DNA Microbiome kit (Qiagen, Hilden, Germany) following the manufacturer’s protocol (*Depletion of Host DNA* protocol).

### Library preparation and sequencing

Twenty-four multi-amplicon libraries were prepared using the QIAseq 16S/ITS Screening Panel kit (Qiagen, Hilden, Germany) as detailed below. These libraries were prepared using 22 urine DNA samples, the DNA extraction negative control and the QIAseq 16S/ITS Smart Control (Qiagen, Hilden, Germany), a synthetic DNA sample used both as positive control for library preparation and sequencing, and as control for the identification of contaminants. DNA concentration was determined using the Qubit dsDNA HS Assay kit and Qubit 2.0 Fluorometer (Thermo Fisher Scientific, Waltham, MA, USA). Next, the fungal taxonomic marker internal transcribed spacer (ITS) and six 16S rRNA amplicons, spanning all nine hypervariable regions (V1V2, V2V3, V3V4, V4V5, V5V7 and V7V9), were amplified by PCR. The ITS region was poorly amplified since we used a DNA extraction protocol which depletes eukaryotic DNA. Sequences originated from the ITS amplicon were therefore discarded. PCR primers and their properties (estimated with OligoCalc (14)) are provided in Table S2. Amplifications were carried out in three independent reactions with primers multiplexed by the manufacturer. For samples in which DNA concentration was ≥0.25 ng/ul, 1 ng of DNA was used as template, and for samples with <0.25 ng/ul, 4 ul of DNA was used. Cycling conditions were: 95 °C for 2 min; 20 cycles of 95 °C for 30 s, 50 °C for 30 s and 72 °C for 2 min; and 72 °C for 7 min. PCR products from the same sample were pooled and purified twice using QIAseq beads (Qiagen, Hilden, Germany). Dual-index barcodes and adapters were added to amplified products through a second-round of PCR using the QIAseq 16S/ITS 96-Index I array (Qiagen, Hilden, Germany). Cycling conditions were: 95 °C for 2 min; 19 cycles of 95 °C for 30 s, 60 °C for 30 s and 72 °C for 2 min; and 72 °C for 7 min. After an additional purification using QIAseq beads, the presence of target sequences was evaluated with the Agilent Bioanalyzer 2100 System using the Agilent DNA 1000 kit (Santa Clara, CA, USA). Finally, we quantified the libraries using the NEBNext^®^ Library Quant Kit for Illumina (New England Biolabs, Ipswich, MA, USA), size-correcting for the average length reported in the Bioanalyzer report considering a 400-700 bp quantification window. Libraries were normalized to 2 nM and sequenced using the MiSeq Reagent Kit v3 (600-cycle) (Illumina, San Diego, CA, USA) following the 2 x 276 bp paired-end read protocol.

### Read processing

Paired-end reads were library demultiplexed and adapters were removed in the Illumina BaseSpace Sequence Hub. Each library was amplicon demultiplexed using cutadapt (v3.4) (15), generating two FASTQ files (with forward or reverse reads) for every library-amplicon combination. FASTQ files from the same amplicon were grouped in QIIME 2 artifacts and processed as independent datasets (hereinafter referred to as amplicon-specific datasets) using QIIME 2 (16).

Using DADA2 (17) (*q2-dada2* QIIME 2 plugin), reads were filtered based on default quality criteria, denoised and truncated (at the first instance of median quality score <30) to remove low quality bases at 3’ ends. Next, paired-end reads were merged using DADA2 to produce amplicon sequence variants (ASVs). Finally, chimeric ASVs were filtered using VSEARCH (18) (*q2-vsearch* QIIME 2 plugin) and the SILVA database (v138) (19) as reference.

### Taxonomic assignment, nomenclature homogenization and contaminant removal

Custom slices of the SILVA database (v138) for each amplicon were generated using RESCRIPt (20) (*q2-rescript* QIIME 2 plugin). Low-quality reference sequences were removed, identical reference sequences were dereplicated and 16S rRNA hypervariable regions were selected using primer sequences from the first-round of PCR as target sequences. Only selected regions within a reasonable length-range (100-600 nt) were kept in the final amplicon-specific databases. To achieve a more accurate taxonomic assignment for each amplicon-specific dataset (21), amplicon-specific taxonomic classifiers trained in amplicon-specific databases were built using the *q2-feature-classifier* QIIME 2 plugin (22). Finally, taxonomic assignment of ASVs was performed using amplicon-specific databases and classifiers.

Assigned taxonomies often contain incomplete information or generic proxies, especially at species level. To homogenize taxonomic nomenclature and to prevent inflation of taxonomic richness at species level, we replaced missing data, generic proxies (terms including “_sp.”, “uncultured”, “metagenome”, or “human_gut”) and ambiguous taxonomic entries (e.g., “phylum: Bacteroidota|Proteobacteria)” by the lowest taxonomic level with complete nomenclature and the corresponding taxon (e.g. “(…) genus: Streptococcus; species: uncultured_bacterium” is replaced by “(…) genus: Streptococcus; species: Genus_Streptococcus”).

Next, we filtered non-bacterial and bacterial contaminants using taxonomic and abundance information. Non-bacterial contaminants were filtered by removing ASVs classified as not being from bacterial origin (taxonomy assigned to mitochondria, chloroplast or unassigned kingdom). Bacterial contaminants were identified using the R package *decontam* (23). Briefly, using the DNA extraction negative control and QIAseq 16S/ITS Smart Control libraries as controls for contaminants, we tested whether each ASV was a contaminant by combining frequency and prevalence *decontam* methods. Due to the limited number of DNA extraction negative control libraries, there was limited statistical power to identify contaminants exclusively from abundance data. Therefore, we evaluated manually if potentially contaminant ASVs (P < 0.25) had been previously described as belonging to human microbiotas by searching the taxon associated with such ASVs at PubMed (search in May 2021). Potentially contaminant ASVs whose taxonomy had not been previously described in urine (namely, *Pelomonas*, which is a known laboratory contaminant (24), *Mycoplasma wenyonii* and *Candidatus Obscuribacter* ASVs) were considered true bacterial contaminants and were removed from all amplicon-specific datasets.

### Sidle-reconstruction of amplicons combinations

The Short MUltiple Reads Framework (SMURF) algorithm (25) as implemented in Sidle (SMURF Implementation Done to acceLerate Efficiency) (26) was used to reconstruct datasets combining all six 16S rRNA amplicons. The *q2-sidle* QIIME 2 plugin was used (as described below) with amplicon-specific datasets after contaminants removal.

For ASVs in each amplicon-specific dataset to have a consistent length (as demanded by SMURF algorithm), ASVs were truncated at 300 nt. Reference sequences in the amplicon-specific databases generated previously were also truncated at 300 nt. For each truncated amplicon-specific database, regional k-mers were aligned (with 5 nt maximum mismatch) and a reconstructed database incorporating all amplicon-specific databases was built. Next, we reconstructed the abundance (0 minimum number of counts) and the taxonomic table incorporating all amplicon-specific datasets. Finally, we removed all libraries classified as defective and homogenized taxonomic nomenclature as previously described. Sidle-reconstructed datasets combining pairs of amplicons were built through an analogous pipeline.

### Microbiota analyses

Amplicon-specific datasets were normalized prior to diversity analyses by Scaling with Ranked Subsampling (27) using the R package *SRS* (28). The number of reads of the library with the lowest number of reads per dataset was used as normalization cutoffs. The normalized amplicon-specific datasets were used to compute taxonomic and ASV richness (where richness is defined as the number of different observed features per dataset), and Faith’s phylogenetic diversity index (29) using the R package *picante* (30). Compositional dissimilarity between samples (beta-diversity) was estimated using either Bray-Curtis or Jaccard (31) indices using the R package *phyloseq* (32).

ASVs were aligned using the R package *DECIPHER* (33) to calculate the entropy per nucleotide for each dataset, and the entropy score was calculated using the R package *Bios2cor* (34).

Genera intersections between datasets were determined using the R package *UpSetR* (35). Taxonomic trees were generated using the R package *metacoder* (36) employing the Reingold-Tilford layout. Only the 32 most abundant taxa were shown when plotting taxa relative abundances (based on minimum relative abundance in at least one sample, which is adjusted for each plot).

Ambiguity was estimated using the abundance output tables generated using Sidle. In these tables, the number of potential 16S rRNA source sequences for each feature is provided. For each dataset, ambiguity was calculated as the sum of the log of the number of potential 16S rRNA source sequences for each feature in the abundance table over the total number of features in the abundance table. To calculate the ambiguity for amplicon-specific datasets (not generated by Sidle), Sidle abundance tables for each amplicon were built as described in the previous section.

The full bioinformatics pipeline and R scripts (37) used for plotting (mainly with the R package *ggplot2* (38)) are available at https://github.com/vitorheidrich/urine-16S-analyses.

## Results

### Sequencing output and taxonomic resolution

We were able to amplify target sequences from 18 out of the 22 (82%) urine samples. In total we generated 20 amplicon libraries spanning all 16S rRNA hypervariable regions for urinary microbiota profiling (18 libraries from urine samples and libraries for the DNA extraction negative control and the QIAseq 16S/ITS Smart Control). A total of 13,638,685 reads were generated from urine sample libraries (median per library: 609,902; range: 383,982-1,489,196), out of which ~59% were short unspecific reads not associated with any of the amplicons of interest (the read length distribution of each amplicon-specific dataset is shown in Figure S1). After amplicon demultiplexing, each amplicon-specific dataset was analyzed in parallel. The total number of reads generated for each amplicon-specific dataset varied between 668,509 (V3V4) and 1,674,525 (V4V5) (Figure 1A; Table S3). After read filtering and removal of contaminants (see Methods), amplicon-specific datasets showed on average a 28% decrease in the number of reads (Figure 1A; Table S3). The number of reads removed at each step in our bioinformatics pipeline is detailed in Table S3.

**Figure 1:**
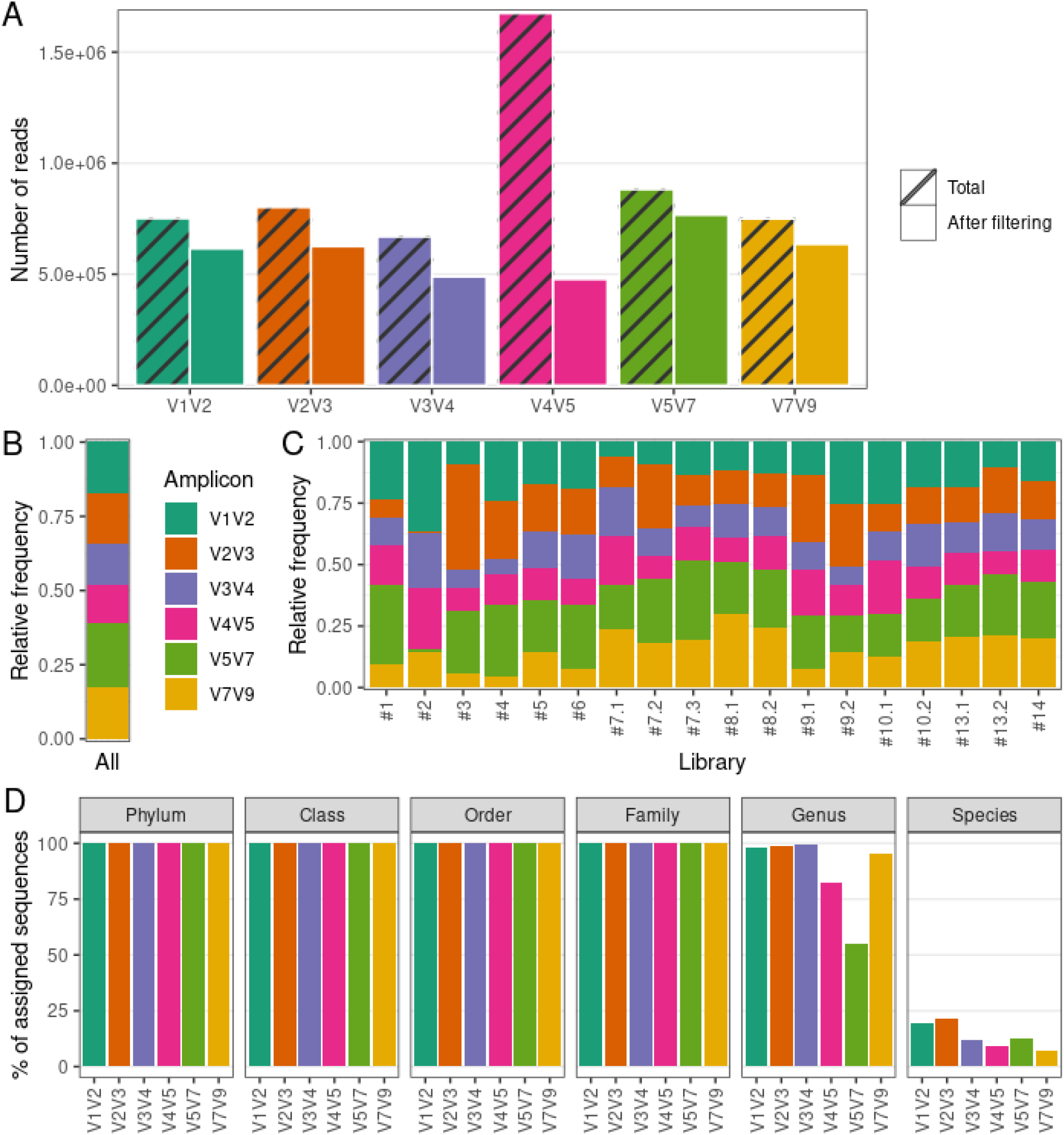
Sequencing output and taxonomic resolution for each 16S rRNA amplicon-specific dataset. (A) Number of reads generated and retained after filtering steps for each amplicon-specific dataset. (B) Relative frequency of reads retained after filtering steps averaged over all libraries for each amplicon-specific dataset. (C) Relative frequency of reads retained after filtering steps per library for each amplicon-specific dataset. (D) Percentage of sequences with assigned taxonomy (per taxonomic level) for each amplicon-specific dataset.

Despite an overall balanced relative abundance of reads for each amplicon-specific dataset (Figure 1B), some libraries presented a disproportionate number of reads for a particular amplicon (Figure 1C). Specifically, libraries #2 and #3 showed a high proportion (>⅓) of V1V2 and V2V3 reads, respectively. We also noted that, despite the high median total number of reads generated for each library (204,433), the extremes varied by orders of magnitude (from 5,366 to 504,852 reads), so that the library with the lowest number of reads (#1) had less than 1,000 reads in 4 out of 6 amplicon-specific datasets. These disparities lead us to remove libraries #1, #2 and #3 from further analyses to prevent the introduction of bias due to low-quality libraries. Finally, we confirmed that the remaining libraries achieved satisfactory sequencing depth by calculating the Good’s coverage (39) (~100% for all samples) and drawing rarefaction curves (Figure S2) for each amplicon-specific dataset.

Within these refined datasets, virtually all sequences in V1V2, V2V3 and V3V4 datasets received a taxonomic assignment up to genus level (Figure 1D). On the other hand, V4V5 and V5V7 showed a marked decrease in the percentage of assigned sequences at genus level, suggesting a lack of taxonomic resolution for relatively abundant taxa. Taxonomic assignment up to species level was rarely achieved, with V1V2 (19.7%) and V2V3 (21.8%) datasets showing the highest percentage of sequences assigned up to species level.

In summary, our results indicate that the protocol used herein is suitable for urinary microbiota characterization, providing enough sequencing depth to assess several amplicons simultaneously. We also confirmed that 16S rRNA hypervariable regions sequencing of urine samples can provide reliable taxonomic information up to genus level. However, taxonomic resolution varies along the 16S rRNA hypervariable regions, with V1V2 and V2V3 achieving the highest taxonomic resolution when considering genus and species levels together.

### Richness across 16S rRNA amplicon-specific datasets

Next, we evaluated ASV and taxonomic (phylum to species level) richness for each amplicon-specific dataset (Figure 2A-B). V1V2 and V3V4 datasets showed the highest ASV richness, while V4V5 and V7V9 presented a markedly lower ASV richness (Figure 2A). There was no correlation between ASV richness per dataset and the median ASV length per dataset (Spearman ⍴ = −0.37, P = 0.47). The ASV length distribution of each amplicon-specific dataset is shown in Figure S3. At phylum and class level, all amplicons showed a remarkably similar richness (Figure 2B), with 6 phyla and 10 classes observed for all datasets, except for the V3V4 dataset (7 phyla and 11 classes). At lower taxonomic levels, differences between amplicons emerged, with the V1V2 dataset showing consistently the highest taxonomic richness from order to species level (Figure 2B). There was no correlation between taxonomic richness per dataset and the median ASV length per dataset (Table S4).

**Figure 2:**
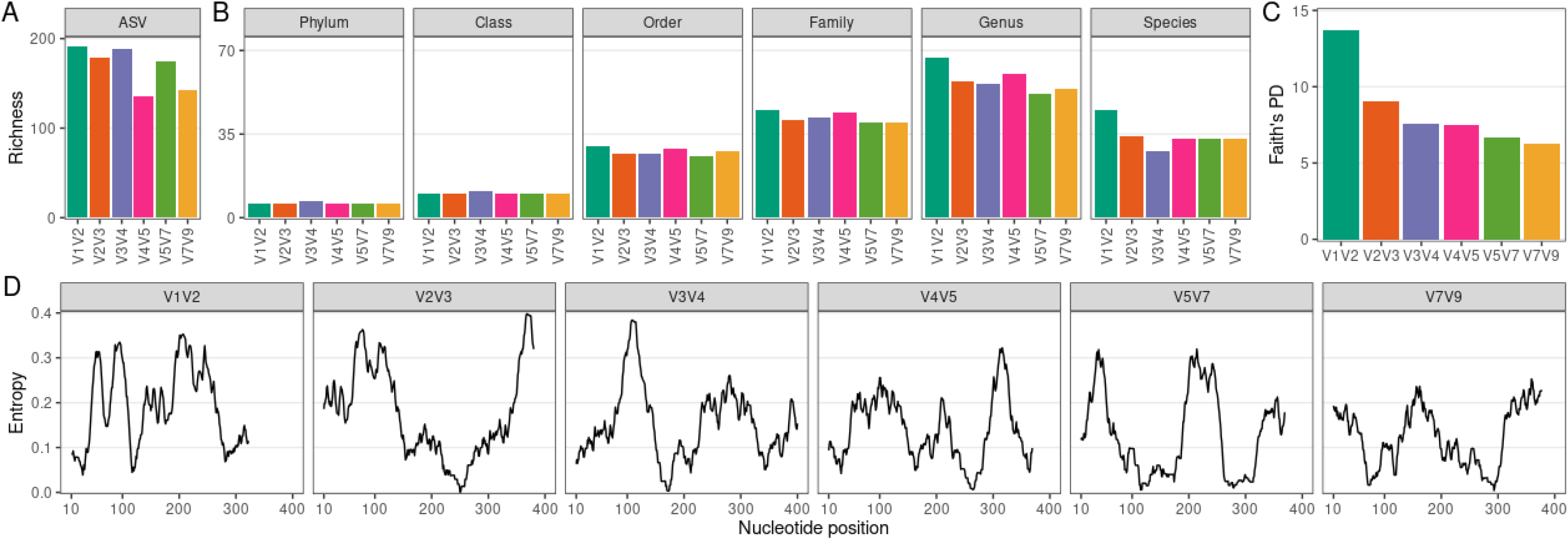
Richness and phylogenetic diversity across 16S rRNA amplicon-specific datasets. (A) Amplicon sequence variant (ASV) richness per amplicon-specific dataset. (B) Taxonomic richness (phylum to species level) per amplicon-specific dataset. (C) Faith’s Phylogenetic Diversity (PD) across amplicon-specific datasets. (D) Sequence variability (entropy) along ASVs nucleotide positions (20-nucleotides rolling average) for each amplicon-specific dataset. Only nucleotide positions up to the median ASV size per amplicon-specific dataset are considered.

As expected from its higher ASV richness, V1V2 showed the highest taxonomic richness at genus level. However, we noticed that ASV richness did not always translate into taxonomic richness. For instance, V3V4 goes from the 2nd to the 4th position when richness was assessed at genus level instead of ASV level, suggesting that part of its ASVs correspond to ASVs phylogenetically close to other ASVs observed in the dataset, which do not contribute to increase taxonomic richness. Indeed, V3V4 ASVs showed a much lower phylogenetic diversity compared to V1V2 ASVs (Figure 2C). In fact, there is a decreasing trend in phylogenetic diversity along the 16S rRNA hypervariable regions, which is in line with the sequence variability (entropy) observed for each amplicon-specific dataset (Figure 2D). There was no correlation between ASV phylogenetic diversity and the median ASV length (Spearman ⍴ = −0.09, P = 0.92).

Together, our results indicate that V1V2 is the most informative 16S rRNA amplicon in terms of taxonomic richness and phylogenetic diversity for urinary microbiota characterization.

### Taxonomic composition across 16S rRNA amplicon-specific datasets

The phyla Actinobacteriota, Bacteroidota, Firmicutes, Fusobacteriota and Proteobacteria were detected in all amplicon-specific datasets. However, some phyla were detected exclusively in a subset of them (Figure S4A). Therefore, we analyzed how taxa detection varied across amplicon-specific datasets at genus level. The full picture of the genera detected in each amplicon-specific dataset is provided in Figure S4B. Taxonomic trees depicting the contribution of each taxon (tree nodes) to the genera detected in each dataset are provided in Figure S5.

When evaluating the intersection of genera present in each amplicon-specific dataset (Figure 3A), we see that 27 genera were detected in all amplicon-specific datasets. The next larger subgroup, composed of 15 genera, comprises genera detected exclusively in the V1V2 dataset. Noteworthy, the V1V2 dataset is the only amplicon-specific dataset without exclusively undetected genera. All other datasets also presented “exclusive” genera, which summed up to 31 genera.

**Figure 3:**
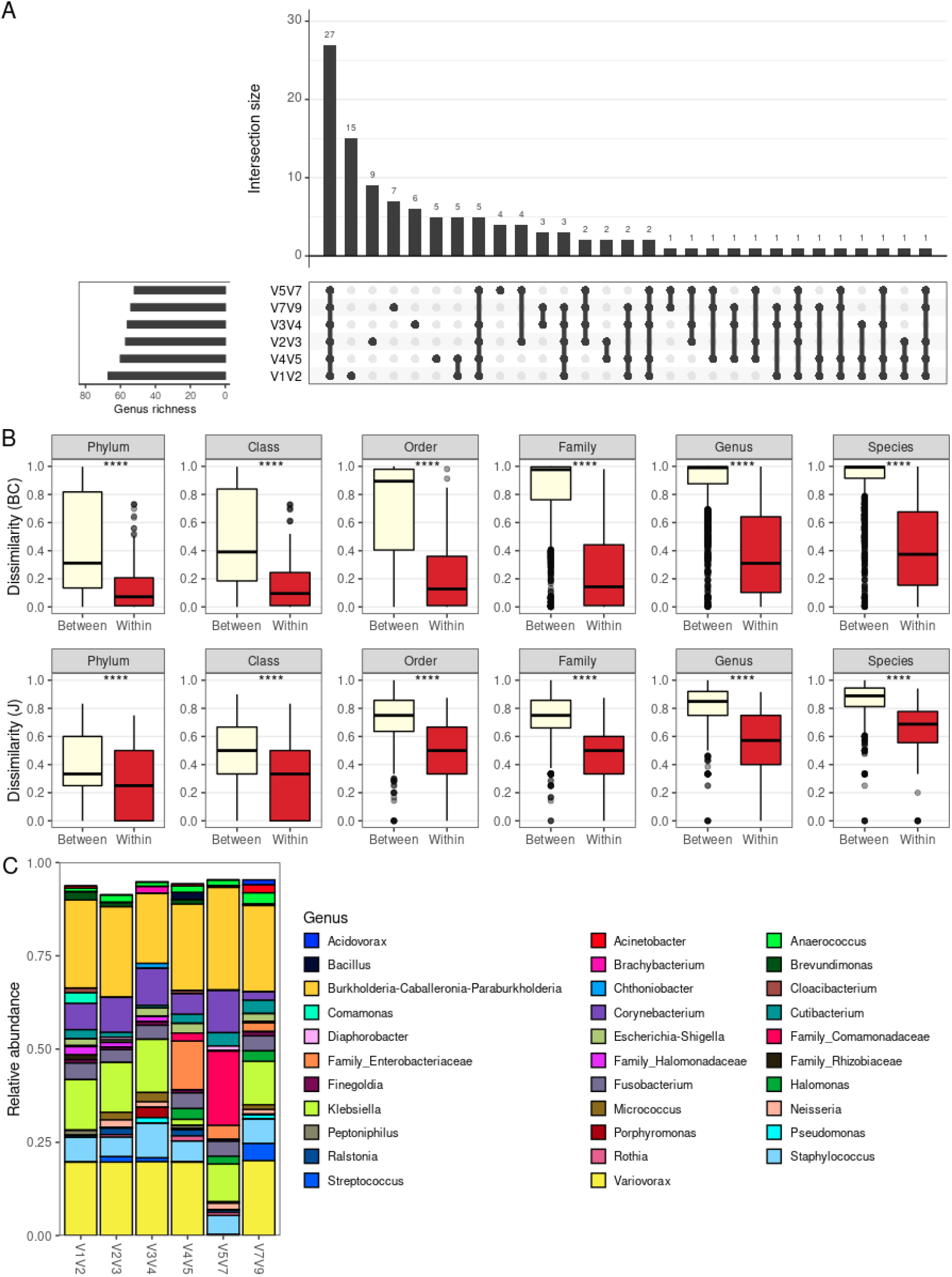
Taxonomic composition across 16S rRNA amplicon-specific datasets. (A) Barplot depicting intersections between the genera detected in each amplicon-specific dataset. Total rIchness at genus level is shown in the lower-left subplot. (B) Boxplot comparing dissimilarities between different libraries and within the same libraries as profiled with different amplicons. Dissimilarity metrics considered are Bray-Curtis (BC) and Jaccard (J). Statistical significance was evaluated by the Mann-Whitney U test. The boxes highlight the median value and cover the 25th and 75th percentiles, with whiskers extending to the more extreme value within 1.5 times the length of the box. (C) Average genera relative abundance per amplicon-specific dataset. ***, P < 0.001.

Due to such substantial differences, we aimed to assess how the choice of amplicon affects taxonomic profiles. To do so, we used beta-diversity analysis to evaluate whether the taxonomic composition of a given sample is similar to itself irrespectively of the amplicon used for characterization (Figure 3B). For all taxonomic levels, using either Bray-Curtis or Jaccard beta-diversity indices, the compositional dissimilarities within samples (same sample profiled with different amplicons) are significantly lower than between samples, suggesting that the choice of amplicons will marginally impact the overall taxonomic compositions, especially at higher taxonomic levels.

The robustness of the taxonomic profile obtained irrespectively of the amplicon of choice can be further contemplated by the similar genera relative abundance profile (averaged over all samples) obtained for each amplicon-specific dataset (Figure 3C). In Figure 3C, there is an apparently disparate average taxonomic composition for V4V5 and V5V7 datasets. However, this is mainly due to loss of taxonomic resolution for some taxa, with ASVs otherwise classified as genera *Variovorax* and *Klebsiella* being only resolved up to family level (Comamonadaceae and Enterobacteriaceae, respectively) in these datasets. This loss of taxonomic resolution is also observed for *Halomonas* ASVs, which were classified as so in V4V5, V5V7 and V7V9 datasets, but as “Family_Halomonadaceae” in the remaining ones. This phenomenon is even more evident when evaluating taxa relative abundances per sample for each dataset at different taxonomic levels (Figure S6), with examples of higher taxonomic resolution at species level (e.g. for *Staphylococcus sp*. in V1V2 and V2V3 datasets).

Despite small variations in taxonomic resolution across amplicon-specific datasets for specific taxa, the overall taxonomic composition of urinary samples is similar independently of the amplicon of choice. Still, each amplicon is able to capture a different subset of the taxa, with V1V2 providing the highest number of exclusively detected genera. These results are in line with the higher genus richness observed for the V1V2 dataset and indicate that V1V2 better captures the actual microbiota composition of urinary samples.

### Comparison with Sidle-reconstructed datasets

We next evaluated how the taxonomic richness and composition differ when considering a single amplicon-specific dataset vs. the Sidle-reconstructed taxa abundance, considering the information for all amplicon-specific datasets simultaneously. This “full” dataset can then serve as a compiled reference for urinary microbiota analysis. We also used Sidle to reconstruct taxa abundances for the following pairs of non-overlapping amplicons: V1V2-V4V5, V1V2-V5V7, V1V2-V7V9, V2V3-V5V7, V2V3-V7V9 and V4V5-V7V9.

As expected, there is a considerable gain in richness in the full dataset, mainly at species level, with 3.9x more species observed in the full dataset when compared to amplicon-specific datasets (Figure 4A). The use of pairs of amplicons also increases richness, but to a lower extent (Figure S7A), with V2V3-V7V9 combination providing the greatest increase in richness at species level (1.9x). This result can be explained by a more complete taxonomic assignment being achieved for a greater proportion of sequences in the full dataset (Figure S7B). Indeed, better taxonomic resolution observed for the Sidle-reconstructed datasets is due to the lower ambiguity (see Methods) in taxonomic assignment (Figure 4B). Noteworthy, the V1V2 dataset shows the lowest ambiguity when comparing only single amplicon-specific datasets.

**Figure 4:**
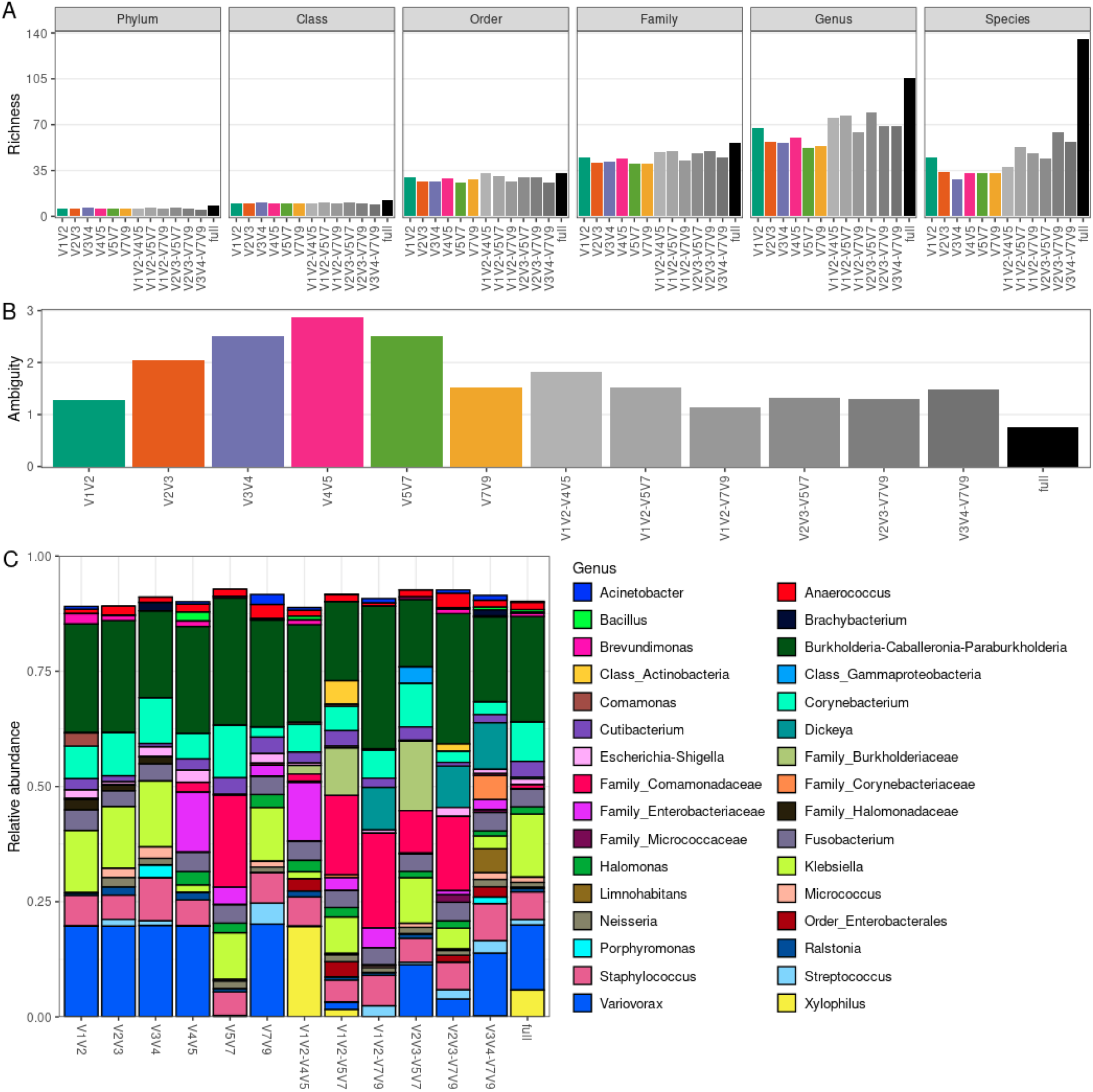
Richness and taxonomic composition of Sidle-reconstructed datasets. (A) Taxonomic richness (phylum to species level) per amplicon-specific or Sidle-reconstructed dataset. (B) Ambiguity in taxonomic assignment per amplicon-specific or Sidle-reconstructed dataset. (C) Average genera relative abundance per amplicon-specific or Sidle-reconstructed dataset.

Once again, the overall taxonomic composition is similar between datasets at genus level (Figure 4C). However, we see cases in which identification at species level was only possible in Sidle-reconstructed datasets (e.g., *Klebsiella pneumoniae* was identified in the full dataset and in most of the pairs of non-overlapping amplicons combinations) (Figure S7C). The taxa relative abundance per sample for the Sidle-reconstructed datasets at different taxonomic levels is provided in Figure S8.

Overall, the combination of amplicons through Sidle increases the taxonomic resolution achievable from 16S rRNA amplicon sequencing. However, the increase of combining pairs of amplicons is modest compared to the full reconstruction using all 16S hypervariable regions, which increases up to 4-fold the number of species detected. Still, this has limited impact in the taxonomic compositions, as evaluated by comparison with the taxonomic profiles generated by single amplicons. Once again, V1V2 stands out as the least ambiguous amplicon for urinary microbiota characterization.

### V1V2 taxonomic composition

Due to the great number of taxa identified in the V1V2 dataset, we next investigated whether these taxa are commonly associated with the urogenital microbiota. In our cohort, six phyla were detected using V1V2 amplicon sequencing: Proteobacteria (72.4% of the sequences), Firmicutes (11.3%), Actinobacteriota (9.6%), Fusobacteriota (4.5%), Bacteroidota (2.3%) and Campilobacterota (<0.1%). All of these phyla have been previously reported in studies using catheterized urine samples (40–42). Only one of such studies reported the overall phyla abundance. The top-three most abundant phyla in Mansour et al. (2020) were Firmicutes, Proteobacteria and Actinobacteriota (42). However, their cohort included females, which are known to have a Firmicutes-enriched urogenital microbiota due to the high abundance of lactobacilli (43). In fact, a study with voided urine specimens from male bladder cancer patients found the same top-three most abundant phyla as described in this study (6).

Next, we examined the 15 genera detected exclusively in the V1V2 dataset. The average relative abundance of these genera varied between <0.001% (*Alkalibacterium* and *Jeotgalibaca*) and 2.9% (*Comamonas*), summing up to ~4% of the bacterial microbiota exclusively detected by V1V2 16S amplicon sequencing (Table S5). Due to the overall low relative abundance of these genera, we excluded the possibility of them being contaminants by searching the literature for the presence of these genera in urine samples. Briefly, 12 out of the 15 (80%) genera exclusively detected in the V1V2 dataset have been previously detected in human samples, and 10 out of 12 (83%) have been associated with urinary infections or detected in urogenital microbiota (Table S5). The three genera that were not previously detected in human microbiotas (*Alkalibacterium*, *Chromohalobacter*, *Salipaludibacillus*) sum up to only <0.01% average relative abundance in the V1V2 dataset. They have been described mainly as environmental high salt tolerant bacteria (44–46), indicating they may indeed represent undetected contamination or taxonomic misclassifications.

Finally, we compared our results with 16S amplicon sequencing-based microbiota studies using catheterized urine samples. In Forster et al. (2020), the urinary microbiota from 34 children with neuropathic bladder was characterized by V4 amplicon sequencing (47). More than 75% of the samples were dominated (relative abundance >30%) by family Enterobacteriaceae members, but the genera involved in this phenomenon could not be determined due to limited taxonomic resolution. We also observed dominance by Enterobacteriaceae members in this cohort (samples #8.1 and #8.2; Figure S6), but because all nine Enterobacteriaceae ASVs in the V1V2 dataset were classified up to genus level (either to *Klebsiella* or *Escherichia-Shigella*), we were able to determine that *Klebsiella sp*. were responsible for this phenomenon. Noteworthy, in V4V5 and V5V7 datasets, family Enterobacteriaceae ASVs could not be classified up to genus level (Figure S6), recapitulating the limited taxonomic resolution for family Enterobacteriaceae observed in the aforementioned study.

Together, these data corroborate that V1V2 amplicon sequencing can provide reliable and richer taxonomic information for microbiota profiling of catheterized urine samples.

## Discussion

Many studies have compared the performance of different sets of 16S rRNA primers for microbiota profiling in different environments (10–13). These studies consistently demonstrated that the choice of the 16S rRNA primer set can significantly influence the analysis of microbiota diversity and composition. Similar studies for urinary microbiota profiling are lacking. As reviewed by Cumpanas et al. (2020) (9), out of 38 urobiome studies, 17 evaluated the V4 and 4 evaluated the V3V4 16S rRNA hypervariable regions. This is probably because these amplicons are commonly used in 16S rRNA amplicon sequencing commercial kits. It is also worth mentioning that some of the early seminal studies were based on V1V3 amplicon sequencing using the Roche 454 platform (2), which allows longer reads. Therefore, up to now library preparation kits and sequencing platforms have heavily influenced the choice of 16S rRNA hypervariable regions used in urinary microbiota profiling studies. Consequently, studies that provide evidence for a more informed choice are urgent.

In this study, we tested the performance of six different 16S rRNA primer sets, spanning all nine hypervariable regions, for microbiota profiling of 22 urine samples collected from male volunteers by transurethral catheterization. We show that V1V2 amplicon sequencing is more suitable for urinary microbiota profiling. We found that V1V2 provides the greatest taxonomic and ASV richness, which translates into a higher number of exclusively detected genera. This result is likely attributed to V1V2 having a higher taxonomic resolution for assessing the taxa commonly present in human urine samples.

We also evaluated combinations of pairs of non-overlapping amplicons, from which we observed only marginal gains in taxonomic richness in comparison with single amplicons. Combining all six amplicons leads to a substantial increase in taxonomic richness at species level, but with little impact on the overall taxonomic compositions, indicating these gains are largely due to low-abundant taxa. Therefore, they may not compensate for the higher costs of sequencing multi-amplicon libraries. Moreover, as amplicon combinations cannot be reconstructed as single sequences, the eventual equivocal association between amplicons may have caused inflation of taxonomic richness by false-positive taxa in Sidle-reconstructed datasets.

We observed huge discrepancies between amplicon-specific datasets when evaluating bacterial compositions by taxa relative abundances. This is mainly because some amplicons presented lower taxonomic resolution for profiling specific clades. Low taxonomic resolution may impact community-wide metrics and preclude the identification of associations between taxa and covariates. Furthermore, low taxonomic resolution may also drastically impact beta-diversity metrics that do not take phylogenetic information into account (e.g., Bray-Curtis).

Amplicon-specific datasets also differed in the set of taxa detected. V1V2 profiling minimized the number of undetected genera, but because all other datasets possessed exclusively detected genera, we conclude that missing a fraction of the urine bacterial richness is inevitable with 16S rRNA amplicon sequencing. Still, low relative abundance taxa drive these observed differences so that analyses will not be harshly influenced by this limitation, except when evaluating beta-diversity with metrics that do not take bacterial evenness into account (e.g., Jaccard).

In this study, removal of contaminants was a key step, since laboratory and reagent contaminants disproportionately affect the microbiota profiling of low bacterial biomass samples (7). However, the method used for contaminant removal has limitations. Since we had low statistical power to detect contaminants exclusively using sequencing data, we validated our findings using information available in the literature. This is questionable because most urobiome studies available did not use strategies to control for contaminants (9), therefore a previous description of a taxon in human urobiomes does not imply it is a true urinary tract-resident microbe. On the other hand, some lists of known reagent and laboratory contaminants are available in the literature (e.g., (24)), but many of the taxa included in such lists are known to be present in human microbiotas. Obviously, these disputes are more frequent when studying less characterized environments. For instance, the genus *Variovorax*, which dominated a few samples in our study, is described as a contaminant by Salter et al. (2014) (24). At the same time, in a contaminant-controlled study, a *Variovorax* strain was identified in the urethra of a non-chlamydial non-gonococcal urethritis patient (48).

Another important limitation of urobiome studies is the lack of information on what are the true microbial members of the human urobiome. Urine samples were not included within the Human Microbiome Project, and due to the lack of an external reference, we focused on comparisons between amplicon-specific datasets. In addition, to partially mitigate the lack of a reference, we also compared amplicon-specific datasets to a bioinformatic reconstruction of the microbial community present in the urine samples using the full set of 16S hypervariable regions. Although further studies with experimentally validated references will be needed to confirm our findings, the results presented can guide methodological decisions in future urobiome studies.

Because genitalia and the urinary tract contain distinct bacterial communities (49), an important variable in urobiome studies is the choice of the sampling method (50). Many urobiome studies evaluate voided urine samples (9), which may contain bacteria from the urethra and genital skin, such that voided urine samples represent the whole urogenital tract. In this study, we evaluated urine samples collected via transurethral catheterization, which reduces the presence of distal urinary tract contaminants compared to voided urine (51). This sampling method, similarly to suprapubic aspiration, allows the specific characterization of the urinary bladder microbiota (52). Even though this was a fundamental consideration to avoid cross-site contamination, further studies will be necessary to evaluate whether our results extend to voided urine specimens. Likewise, since we included only male volunteers in this study, further studies including samples from females are desired to test whether our results can be extrapolated to the female urobiome.

In conclusion, similarly to other reports of primer bias in microbiota studies, we provided strong evidence that V1V2 is the most suitable 16S rRNA amplicon for the characterization of catheterized urine samples microbiotas. To our knowledge, this is the first study to address this question by systematically analyzing all 16S hypervariable regions. This is true not only for catheter-derived urine samples, but actually for any kind of urine sample. We believe that our results might help other researchers make an informed decision about which 16S rRNA hypervariable regions to use for urobiome analysis.

## Data Availability Statement

Raw sequencing data was deposited in the European Nucleotide Archive (ENA) at EMBL-EBI under accession number PRJEB49145 (https://ebi.ac.uk/ena/browser/view/PRJEB49145).

## Ethics Statement

This study was approved by the Ethics Committee of Hospital Sírio-Libanês (#HSL 2018-72). All volunteers provided informed consent to participate before sample collection.

## Author Contributions

Conceptualization and study design: VH, AM, DB, DJ, MA, and AC. Volunteer recruitment and clinical evaluation: AM, DB, DJ, and MA. Samples preparation and sequencing: VH and LI. Bioinformatics and statistical analyses: VH. Writing original draft: VH and AC. Reviewing and editing the manuscript: PA, FB, and AM. Supervision: AC. All authors contributed to the article and approved the submitted version.

