## Supplementary Material for "Choice of 16S ribosomal RNA primers impacts urinary microbiota profiling"

### 1. Supplementary Figures and Tables

#### 1.1. Supplementary Figures

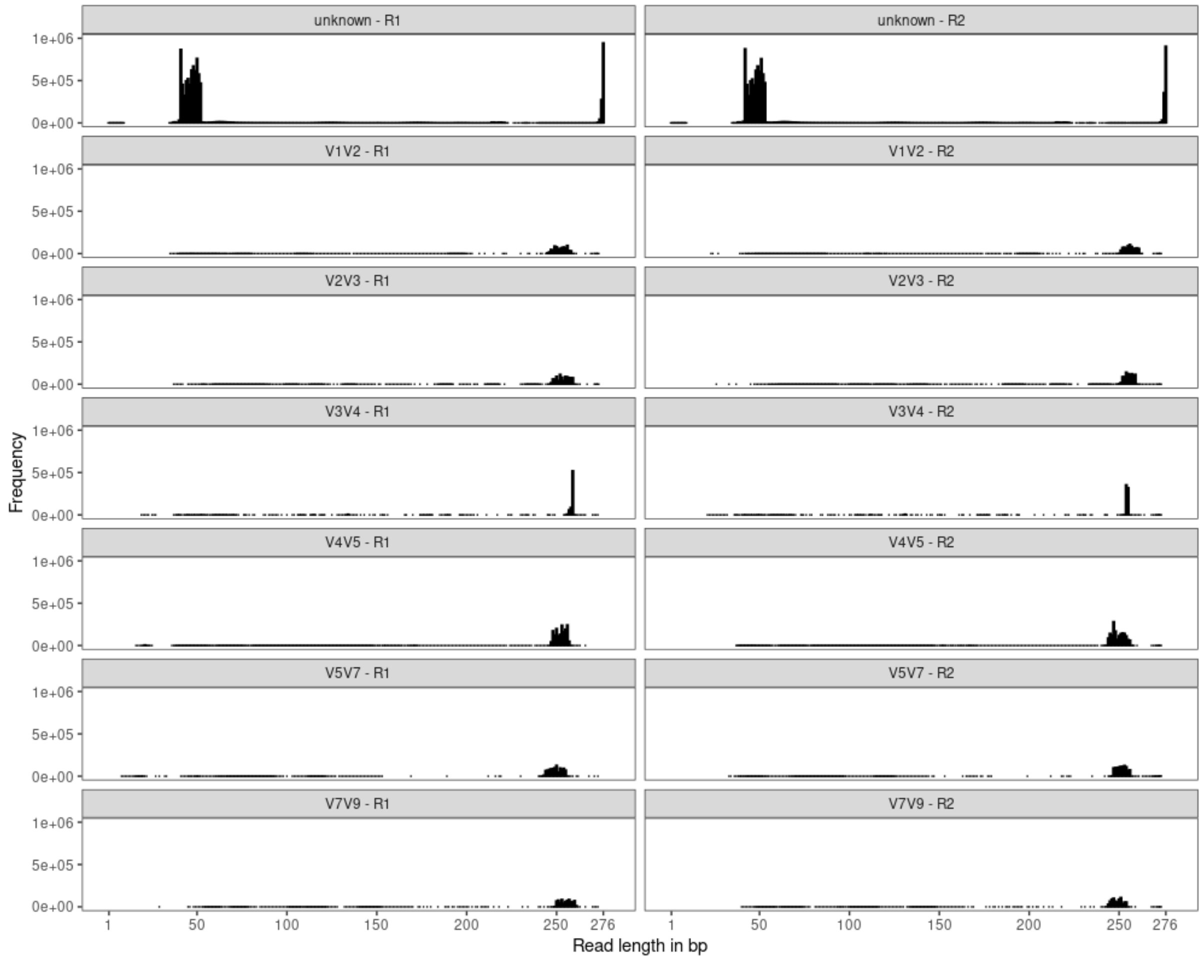


**Supplementary Figure 1: Read lengths for each 16S rRNA amplicon-specific dataset.** Histograms with distribution of read lengths in bp. Read lengths for sequences not associated with any of the amplicons of interest (unknown) are also shown. R1, forward reads; R2, reverse reads.

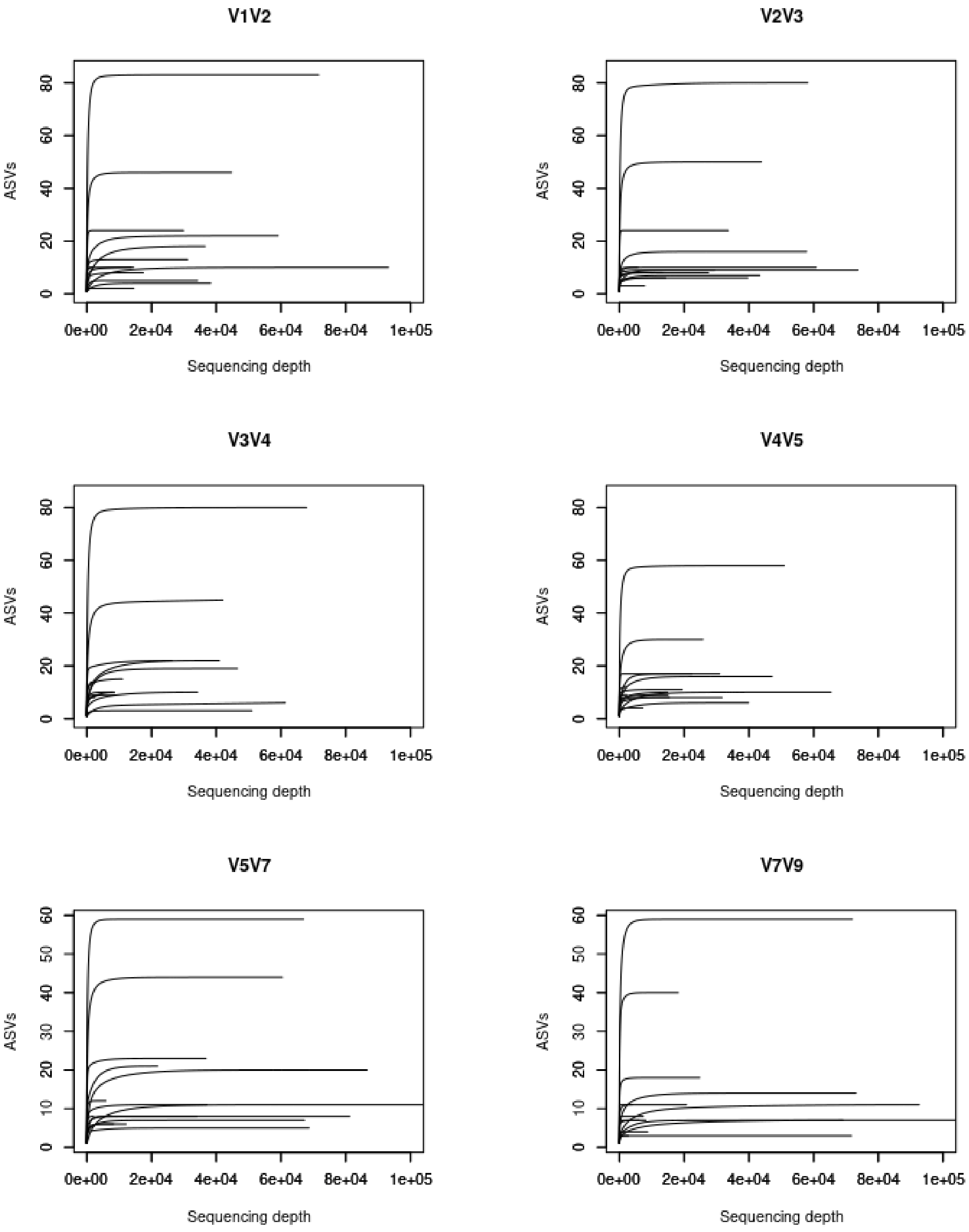


**Supplementary Figure 2: Amplicon sequence variants (ASVs) rarefaction curves for each 16S rRNA amplicon-specific dataset.** Each curve depicts a library. Reads were selected by random subsampling without replacement at incremental steps of 50 reads. Plots were truncated at sequencing depth equals 100,000 reads.

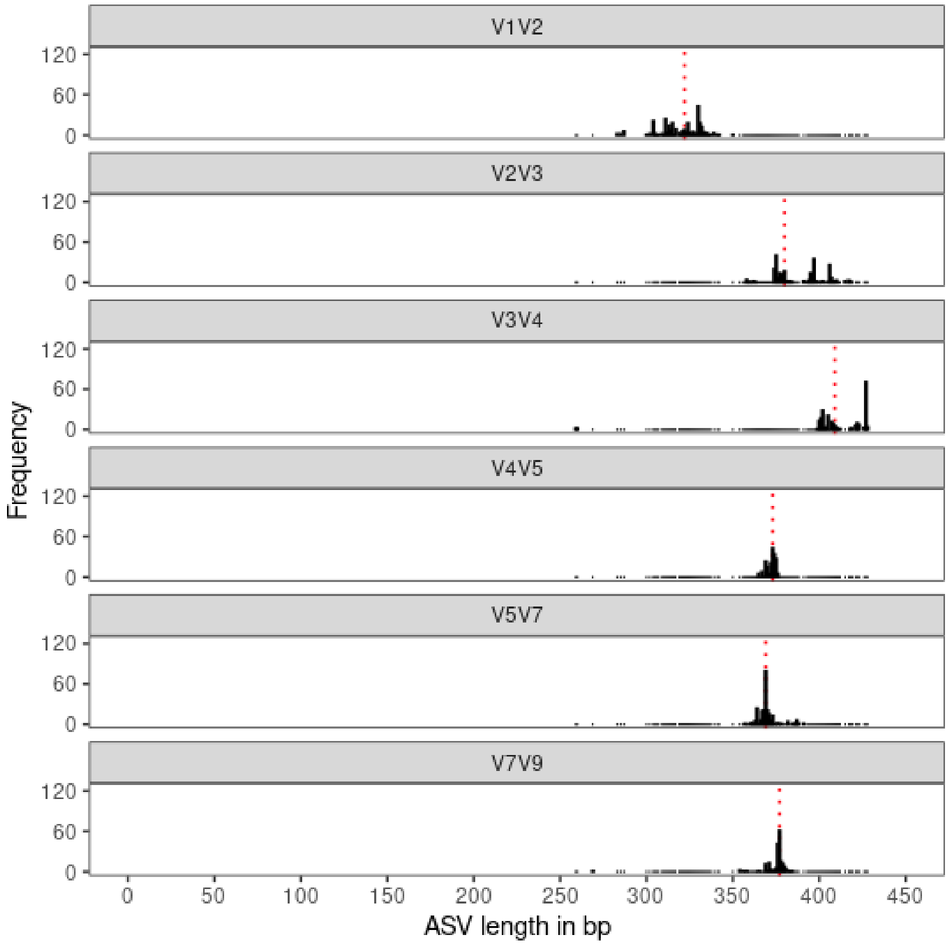


**Supplementary Figure 3: Amplicon sequence variant (ASV) lengths for each 16S rRNA amplicon-specific dataset.** Histograms with distribution of ASV lengths in bp. The median ASV size per amplicon-specific dataset is indicated by a red dotted line.

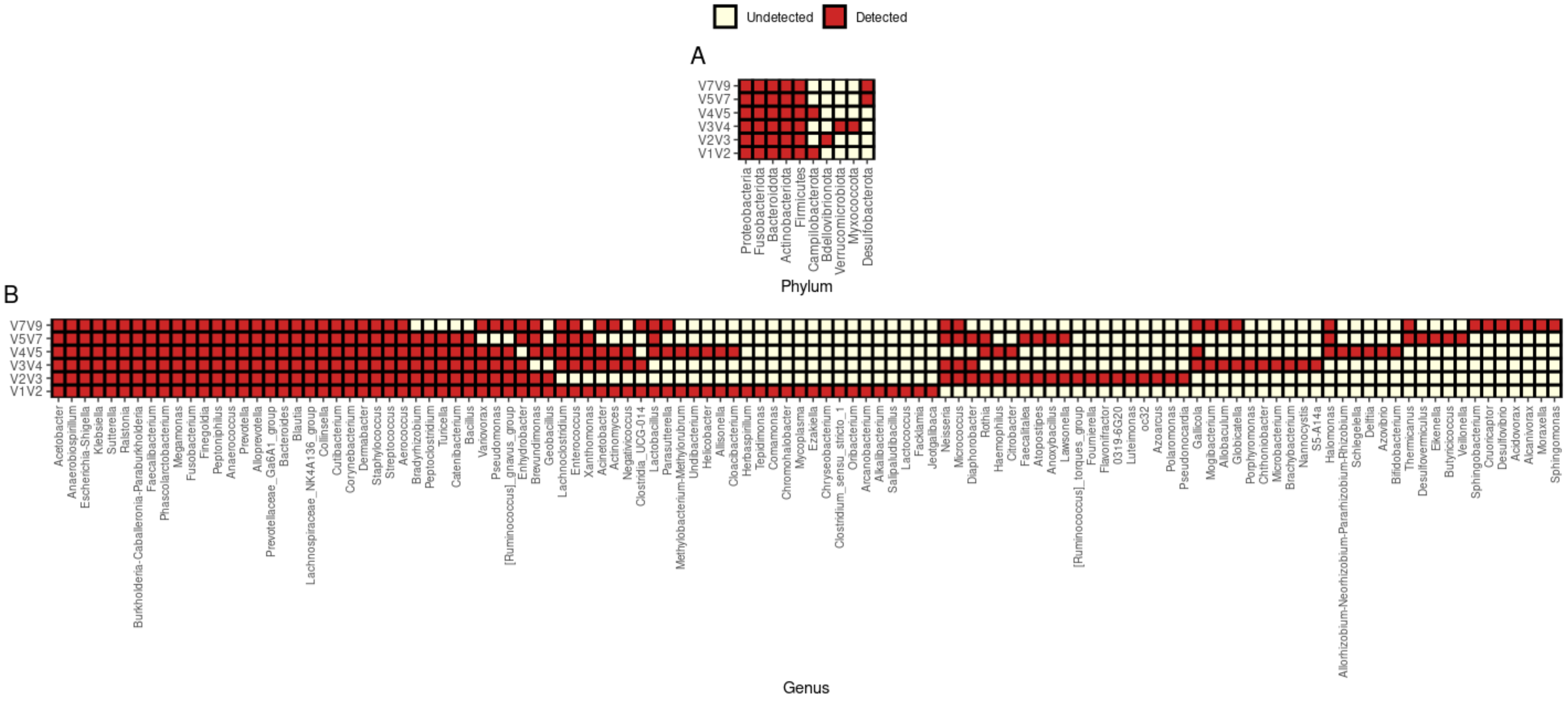


**Supplementary Figure 4: Phyla and genera detected in each 16S rRNA amplicon-specific dataset.** (A) Heatmap depicting the phyla detected in each amplicon-specific dataset. (B) Heatmap depicting the genera detected in each amplicon-specific dataset. Taxa are sorted based on the number of amplicon-specific datasets in which they were detected.

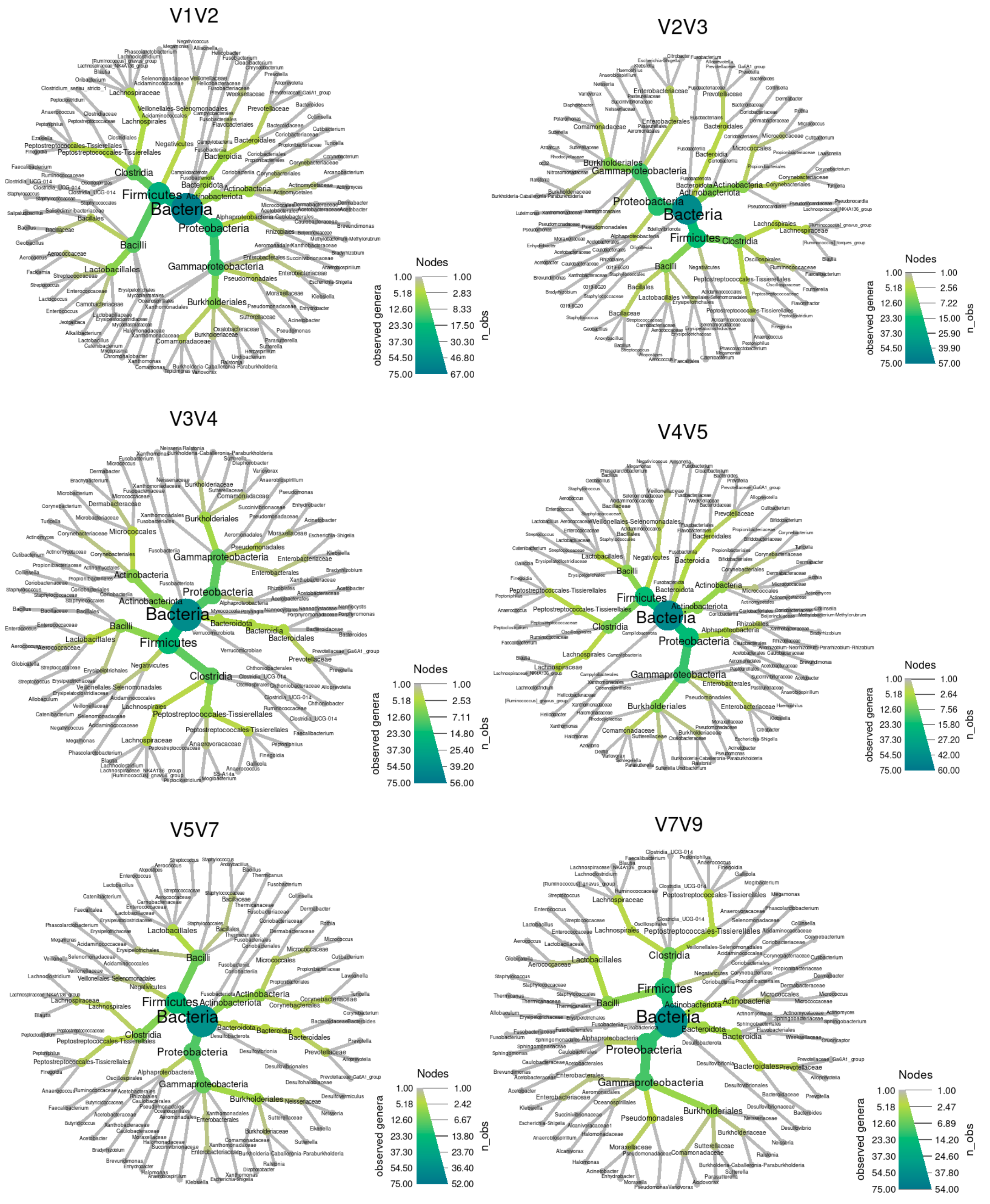


**Supplementary Figure 5: Taxonomic tree of each 16S rRNA amplicon-specific dataset.** Taxa detected in each amplicon-specific dataset are shown up to genus level (tree leaves). Node sizes and color scheme represent the number of leaves (number of genera) present in the subtree of the nodes. The Reingold-Tilford layout was used.

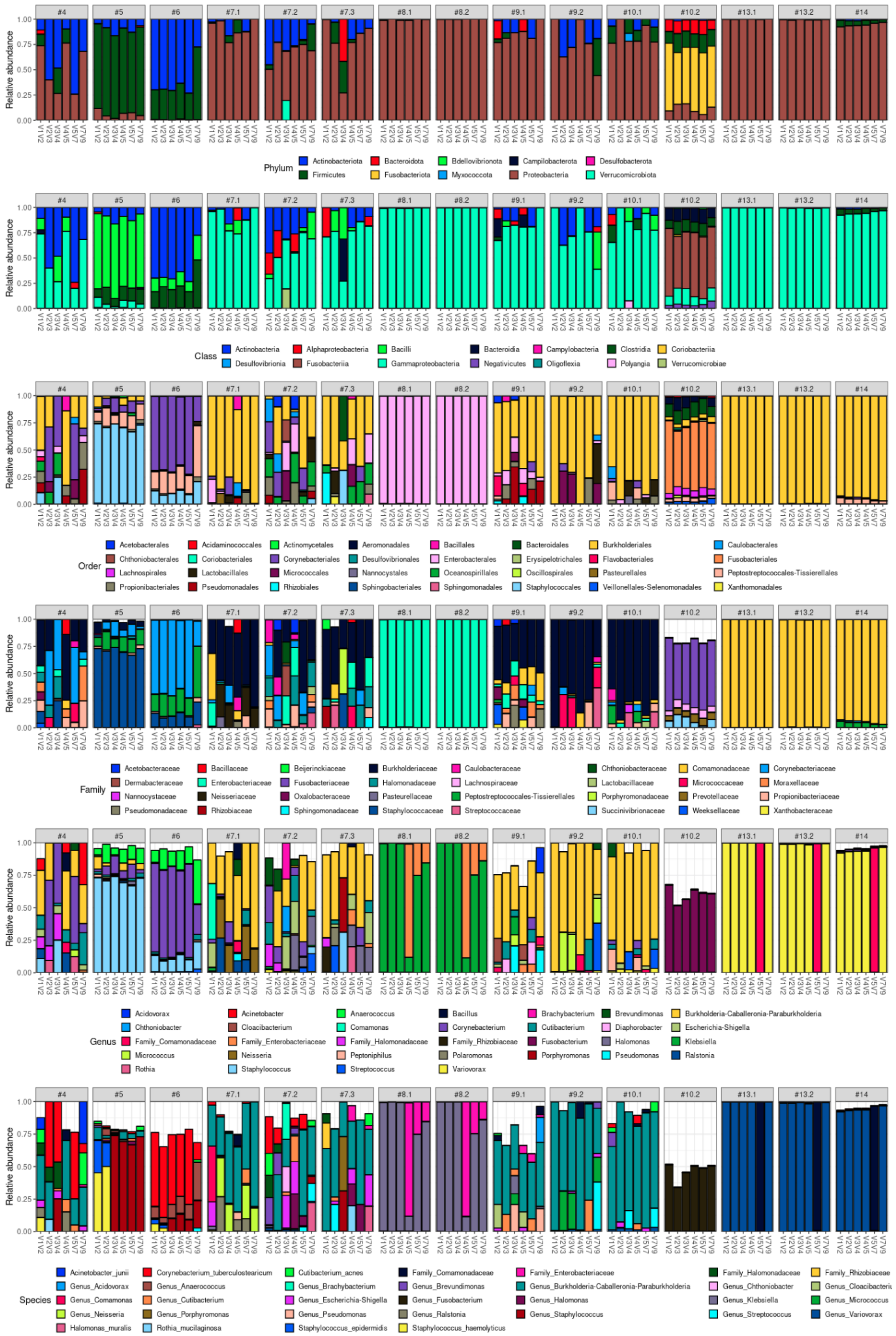


**Supplementary Figure 6: Taxa (phylum to species level) relative abundances for each library as profiled by different 16S rRNA amplicons.**

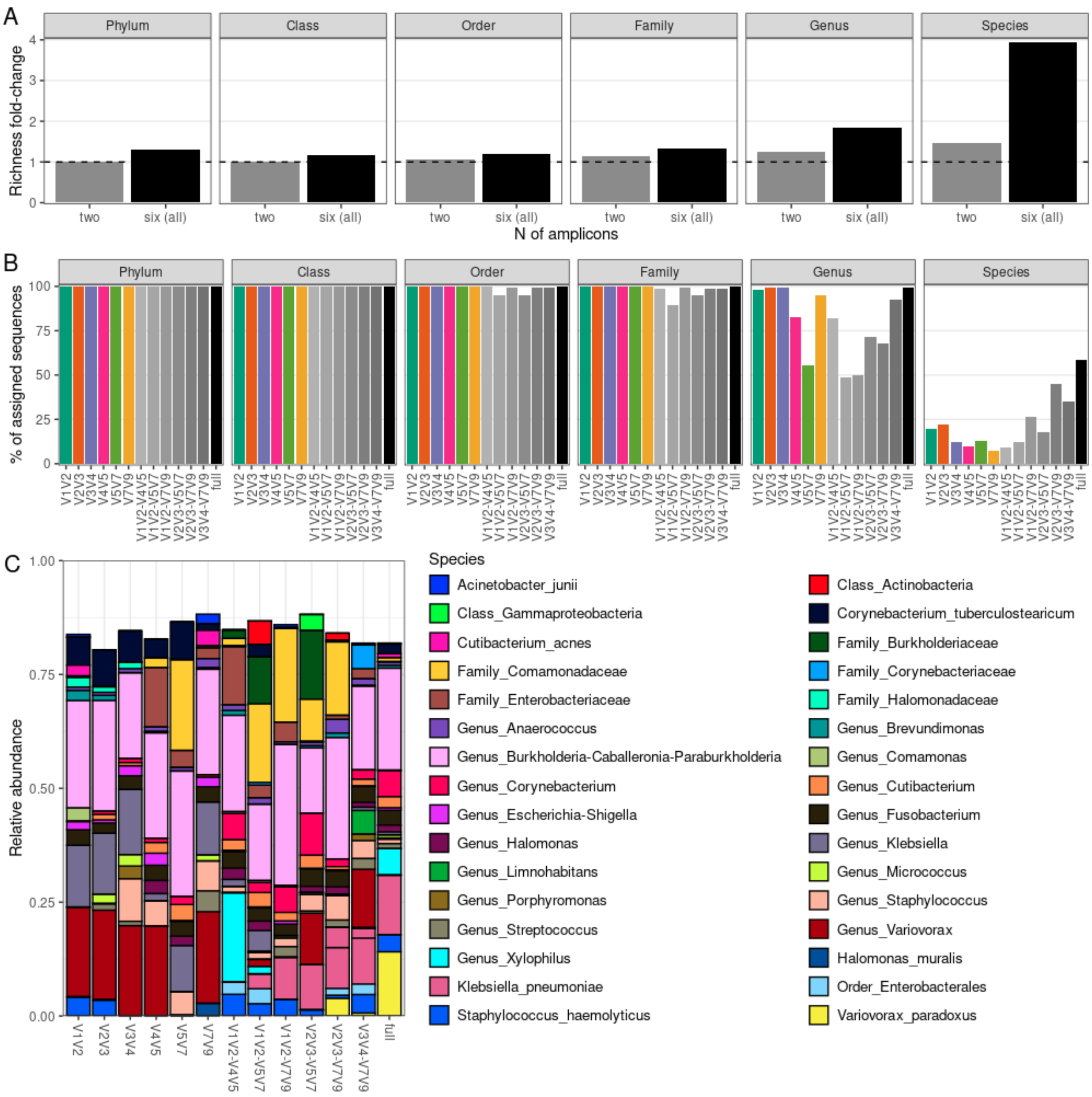


**Supplementary Figure 7: Richness and taxonomic composition of Sidle-reconstructed datasets.** (A) Fold-increase in taxonomic richness of combinations of two (averaged over all pairs) or six amplicons (as built with Sidle) in comparison with the average taxonomic richness found across amplicon-specific datasets. (B) Percentage of sequences with assigned taxonomy (per taxonomic level) for each amplicon-specific or Sidle-reconstructed dataset (C) Average species relative abundance per amplicon-specific or Sidle-reconstructed dataset.

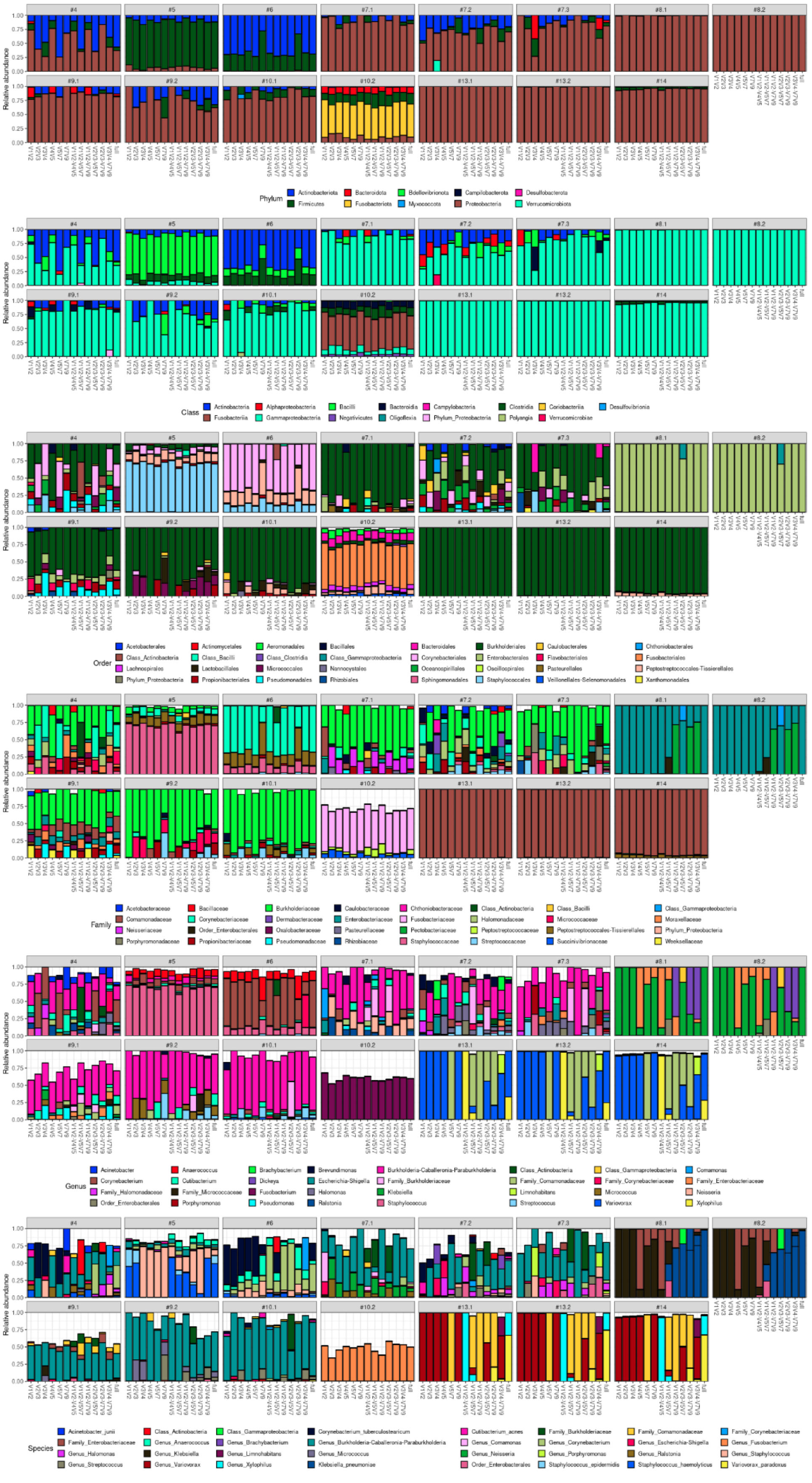

**Supplementary Figure 8: Taxa (phylum to species level) relative abundances for each library as profiled by different 16S rRNA amplicons and Sidle-reconstructed combinations.**

### 1.2. Supplementary Tables

| Patient | Condition | Samples |
|---|---|---|
| #1 | BPH | #1 |
| #2 | BPH | #2 |
| #3 | BPH | #3 |
| #4 | BPH | #4 |
| #5 | BPH | #5 |
| #6 | BPH | #6 |
| #7 | NMIBC | #7.1, #7.2, #7.3 |
| #8 | NMIBC | #8.1, #8.2 |
| #9 | NMIBC | #9.1, #9.2 |
| #10 | NMIBC | #10.1, #10.2 |
| #11 | NMIBC | #11.1*, #11.2* |
| #12 | NMIBC | #12.1*, #12.2* |
| #13 | NMIBC | #13.1, #13.2 |
| #14 | BPH | #14 |
| SC | - | #15 |
| NC | - | #16 |

**Supplementary Table 1: Patients and samples included in the study.** Samples not sequenced due to low target sequence concentration are indicated with an asterisk. NMIBC patients had two or three samples collected longitudinally during the treatment course. BPH, benign prostatic hyperplasia; NMIBC, non-muscle invasive bladder cancer; SC, Smart Control; NC, DNA extraction negative control.

| Amplicon | Primers F | Primers R | Length (F/R) | %GC (F/R) | DI (F/R) |
|---|---|---|---|---|---|
| ITS | CTTGGTCATTTAGAGGAAGTAA | GCTGCGTTCTTCATCGATGC | 22/20 | 36/55 | 0/0 |
| V1V2 | AGRGTTTGATYMTGGCTC | CTGCTGCCTYCCGTA | 18/15 | 39-56/60-67 | 3/1 |
| V2V3 | GGCGNACGGGTGAGTAA | WTTACCGCGGCTGCTGG | 17/17 | 59-65/65 | 3/1 |
| V3V4 | CCTACGGGNGGCWGCAG | GACTACHVGGGTATCTAATCC | 17/21 | 71-76/43-52 | 4/4 |
| V4V5 | GTGYCAGCMGCCGCGGTAA | CCGYCAATTYMTTTRAGTT | 19/19 | 63-74/26-47 | 2/4 |
| V5V7 | GGATTAGATACCCBRGTAGTC | ACGTCRTCCCCDCCTTCCTC | 21/20 | 43-52/60-70 | 3/3 |
| V7V9 | YAACGAGCGMRACCC | TACGGYTACCTTGTTAYGACTT | 15/22 | 53-73/36-45 | 3/2 |

**Supplementary Table 2: List of primers used in the first-round of PCR.** The GC content percentage (%GC) is shown as a range when primers contain degenerate bases. The degeneracy index (DI) of each primer is calculated as the sum of the number of alternate base-pairings for each degenerate base. F, forward primers; R, reverse primers.

| Dataset | Total reads | Median length (F/R) | 25th Q (F/R) | 50th Q (F/R) | 75th Q (F/R) | Q-filtered | Denoised | Merged | Non chimeras I | Non chimeras II | Bacterial | Non contaminants | % Total reads decrease | Median ASV length |
|---|---|---|---|---|---|---|---|---|---|---|---|---|---|---|
| V1V2 | 750,924 | 252/256 | 36/17 | 38/25 | 38/32 | 694,464 | 693,540 | 684,542 | 667,414 | 628,279 | 623,703 | 614,973 | 18,1 | 322 |
| V2V3 | 801,072 | 253/255 | 37/17 | 38/25 | 38/32 | 716,385 | 715,424 | 710,670 | 673,719 | 647,648 | 641,088 | 624,939 | 22,0 | 380 |
| V3V4 | 668,509 | 259/254 | 34/19 | 38/27 | 38/33 | 575,366 | 574,457 | 569,210 | 552,435 | 552,012 | 496,112 | 488,560 | 26,9 | 409 |
| V4V5 | 1,674,525 | 253/249 | 35/15 | 38/23 | 38/30 | 1,491,565 | 1,490,588 | 616,615 | 614,176 | 613,575 | 484,693 | 477,043 | 71,5 | 373 |
| V5V7 | 881,612 | 250/251 | 38/16 | 38/24 | 38/30 | 796,138 | 794,606 | 787,603 | 775,057 | 768,817 | 768,676 | 766,525 | 13,1 | 369 |
| V7V9 | 749,534 | 255/249 | 37/17 | 38/25 | 38/32 | 673,733 | 672,883 | 668,883 | 663,163 | 660,085 | 655,224 | 634,974 | 15,3 | 377 |
| Total | 5,526,176 | | | | | 4,947,651 | 4,941,498 | 4,037,523 | 3,945,964 | 3,870,416 | 3,669,496 | 3,607,014 | | |

**Supplementary Table 3: Reads generated per amplicon-specific dataset.** Metrics on the raw read data and the number of reads retained after each pipeline step are indicated. Only urine sample libraries are considered. Q-score (Q) percentiles are based on the position defined by the last base of the median read. Non-chimeras I and II refers, respectively, to the *de novo* chimera-filtering step within DADA2 and to the *reference-based* chimera-filtering step with VSEARCH. F, forward reads; R, reverse reads; Q-filtered, reads retained after quality-filtering.

| Taxonomic rank | Spearman $\rho$ | P-value |
|---|---|---|
| Phylum | 0,65 | 0,16 |
| Class | 0,65 | 0,16 |
| Order | -0,41 | 0,42 |
| Family | -0,26 | 0,62 |
| Genus | -0,26 | 0,66 |
| Species | -0,58 | 0,23 |

**Supplementary Table 4: Spearman correlation between taxonomic richness and the median ASV length per 16S rRNA amplicon-specific dataset.**

| Genus | Average RA (%) | Presence in urine | Presence in humans |
|---|---|---|---|
| *Comamonas* | 2,86 | Yes (1) | - |
| *Herbaspirillum* | 0,50 | Yes (2) | - |
| *Chryseobacterium* | 0,27 | Yes (3) | - |
| *Ezakiella* | 0,22 | Yes (4) | - |
| *Facklamia* | 0,08 | Yes (5) | - |
| *Mycoplasma* | 0,04 | Yes (6) | - |
| *Oribacterium* | 0,01 | No | Yes (7) |
| *Salipaludibacillus* | <0,01 | No | No |
| *Chromohalobacter* | <0,01 | No | No |
| *Tepidimonas* | <0,01 | Yes (8) | - |
| *Clostridium_sensu_stricto_1* | <0,01 | Yes (9) | - |
| *Arcanobacterium* | <0,01 | Yes (10) | - |
| *Lactococcus* | <0,01 | Yes (11) | - |
| *Jeotgalibaca* | <0,001 | No | Yes (12) |
| *Alkalibacterium* | <0,001 | No | No |
| Sum | 3,99 | - | - |

**Supplementary Table 5: List of genera detected exclusively in the V1V2 dataset.** The average relative abundance (RA) over all samples is shown for each genus. References on the presence of these genera in urine samples or humans are provided.

7. Sizova MV, Muller PA, Stancyk D, Panikov NS, Mandalakis M, Hazen A, Hohmann T, Doerfert SN, Fowle W, Earl AM, et al. Oribacterium parvum sp. nov. and Oribacterium asaccharolyticum sp. nov., obligately anaerobic bacteria from the human oral cavity, and emended description of the genus Oribacterium. *Int J Syst Evol Microbiol* 64:2642–2649. doi: 10.1099/ijs.0.060988-0
